# *MIR822* modulates monosporic female gametogenesis through an ARGONAUTE9-dependent pathway in *Arabidopsis thaliana*

**DOI:** 10.1101/2021.10.18.464879

**Authors:** Andrea Tovar-Aguilar, Daniel Grimanelli, Gerardo Acosta-García, Jean-Philippe Vielle-Calzada, Jesús Agustín Badillo-Corona, Noé Durán-Figueroa

## Abstract

In the ovule of flowering plants, the establishment of the haploid generation occurs when a somatic subepidermal cell specified as the gametophytic precursor differentiates into a Megaspore Mother Cell (MMC) and initiates meiosis. As most flowering plants, *Arabidopsis thaliana* (Arabidopsis) undergoes a monosporic type of gametogenesis as three meiotically derived cells degenerate without further division, and a single one – the functional megaspore (FM) - divides mitotically to form the female gametophyte. The genetic basis and molecular mechanisms that control monosporic gametogenesis remain largely unknown. In Arabidopsis, ARGONAUTE proteins are involved the control of megasporogenesis. In particular, mutations in ARGONAUTE9 (AGO9) lead to the ectopic differentiation of gametic precursors that can give rise to apomeiotically derived female gametophytes. Here, we show that Arabidopsis plants carrying loss-of-function mutations in the AGO9-interacting microRNA miR822a give rise to extranumerary surviving megaspores that acquire a FM identity and divide without giving rise to differentiated female gametophytes. The overexpression of three miR822a target genes encoding Cysteine/Histidine-Rich C1 domain proteins (At5g02350, At5g02330 and At2g13900) results in defects equivalent to those found in *mir822* plants. All three miR822a targets are overexpressed in *ago9* mutant ovules, confirming that miR822a acts through an AGO9-dependent pathway to negatively regulate Cysteine/Histidine-Rich C1 domain proteins and restricts the survival of meiotically derived cells to a single megaspore. Our results identify a microRNA-dependent mechanism that is involved in the control of megaspore degeneration and the most prevalent form of female gametogenesis in flowering plants.

## INTRODUCTION

The life cycle of flowering plants comprises two major stages, the diploid sporophytic and the haploid gametophytic generation. During the sporophytic stage, plants develop and give rise to a mature plant or sporophyte. The female gametophytic phase, which comprises megasporogenesis and megagametogenesis, takes place inside of the female reproductive organs. Megasporogenesis comprises three major events: the specification and differentiation of the Megaspore Mother Cell (MMC), meiosis, and the selection of a single haploid product, the functional megaspore (FM) that will subsequently give rise to the female gametophyte during megagametogenesis (Drews and Koltunow, 2011). In *Arabidopsis thaliana* (Arabidopsis), the formation of the MMC initiates with the enlargement of a single sub-epidermal cell that undergoes meiosis and gives rise to four haploid megaspores, one of which survives and differentiates as the FM. This monosporic pattern of megaspore formation and survival occurs in more than 70% of flowering plants analyzed to date (Huang and Russell, 1992, Haig, 2020). Then, during megagametogenesis, the FM divides mitotically three times without cytokinesis. Cellularization of the resulting eight nuclear *syncytium* gives rise to a differentiated female gametophyte composed of seven cells: three antipodal cells, a binucleated central cell, the egg cell, and two synergid cells. Fertilization of both the egg and central cells triggers a seed developmental program that gives rise to the embryo and endosperm, respectively, and culminates with the formation of a mature seed (Drews and Koltunow, 2011).

Although most flowering plants undergo monospory, there are many species in which more than one meiotically derived nucleus is incorporated into female gametogenesis, suggesting that key steps of megasporogenesis including megaspore death or survival, are controlled by dynamic and variable developmental programs (Schmidt *et al*., 2015, Pinto *et al*., 2019). In most species that follow a monosporic pattern of development, the meiotic nuclear divisions are simultaneous with cytokinesis; however, in other species such as Arabidopsis, nuclear divisions precede cytokinesis (Bajon *et al*., 1999). A tetrad of megaspore nuclei is formed before the deposition of callosic cell walls covering the three dying but not the functional megaspore, suggesting that cell death is - in those cases - a consequence of callose-dependent physical isolation (Rodkiewicz, 1970, Webb and Gunning, 1990). In other species such as *Alisma*, cytokinesis fails after meiosis II, giving rise to two haploid cells, each containing two haploid nuclei. While one of the binucleated cells dies without further differentiation, the second one directly incorporates both of its nuclei into a developing two-nuclear female gametophyte. Finally, in genera such as *Drusa*, cytokinesis is absent after meiosis II, and all four meiotically derived nuclei are incorporated into a four nuclear female gametophyte (Webb and Gunning, 1990, Huang and Russell, 1992, Haig, 2020).

The genetic basis and molecular mechanisms controlling the formation, death, or survival of megaspores remain largely unknown. Using ovules of *Medicago sativa*, some have suggested that programmed cell death (PCD) is the cause of megaspore degeneration (Citterio *et al*., 2005, Drews and Koltunow, 2011). Only a few genes involved in the specification of the FM have been identified. In Arabidopsis, in septuple mutants of *INHIBITORS OF CYCLIN-DEPENDENT KINASES (ICK/KRP)* genes (*ick1 ick2 ick3 ick4 ick5 ick6 ick7*), the number and position of surviving megaspores is variable, indicating that the signals determining survival of megaspore are affected. Nonetheless, these genes act both during MMC specification and meiosis (Cao *et al*., 2018). In addition, three other groups of genes involved in the selection of the Arabidopsis FM have been identified. Overexpression of *ARABINOGALACTAN PROTEIN18 (AGP18)* promotes positive selection of viable megaspores (Demesa-Arevalo and Vielle-Calzada, 2013). By contrast, the specification of the FM is lost in triple mutants of *ARABIDOPSIS HISTIDINE KINASE (ahk2– 7 ahk3–3 cre1–12)* receptor genes (Cheng *et al*., 2013). And in *antikevorkian* (*akv)*, a mutant for which the molecular lesion remains to be determined, extra-numerary survival megaspores can give rise to abnormal female gametophytes (Yang and Sundaresan, 2000).

MicroRNAs (miRNAs) are 21-22 nucleotide (nt) small non-coding RNAs that, together with ARGONAUTE (AGO) proteins, regulate gene expression to control diverse developmental programs in angiosperms (Liu *et al*., 2018). Briefly, the miRNAs are formed from a primary transcript that is synthesized by RNA polymerase II, the resulting non-coding single strand transcript can form a secondary structure hairpin that is usually processed by DICER-LIKE RNAses that generate double-stranded RNA of 21-22 nt in length. The AGO proteins can bind to one of the single-stranded molecules of the RNA duplex and target complementary mRNA sites to either suppress transcription or inhibit translation of the corresponding protein (Armenta-Medina and Gillmor, 2019). In Arabidopsis, miRNAs have been well characterized during vegetative development; however, their role during female gametogenesis remain poorly understood. Yet, genes implicated in the biogenesis of small non-coding RNAs have been linked to early ovule development; for example, the control of cell specification during megasporogenesis is dependent on the RNA-directed DNA methylation (RdDM) pathway, that controls female gamete formation by restricting the specification of pre-meiotic precursors through a silencing mechanism that involves the activity of sRNAs (Duran-Figueroa and Vielle-Calzada, 2010, Olmedo-Monfil *et al*., 2010). Dominant mutations in known genes of the RdDM pathway, including ARGONAUTE9 protein, lead to differentiation of multiple female gametic cells that can initiate gametogenesis without undergoing meiosis by a mechanism reminiscent of apospory (Hernandez-Lagana *et al*., 2016). Although most AGO9 interactors are 24-nt in length and derived from TEs (Duran-Figueroa and Vielle-Calzada, 2010), AGO9 protein can also bind to 21-22 nt miRNAs (Havecker *et al*., 2010, Olmedo-Monfil *et al*., 2010). While several TEs are activated in the egg apparatus in *ago9*mutant alleles, the role of AGO9 miRNA interactors is still unknown.

Here, we analyzed the expression pattern of seven *MIR* genes encoding miRNAs that interact with AGO9: *MIR161, MIR390, MIR858, MIR867, MIR403, MIR158* and *MIR822*. We observed that only *MIR822* showed a defined and specific expression in developing ovules. We genetically analyzed two Arabidopsis *MIR822* null alleles and showed that developing ovules in homozygous mutant lines exhibit several meiotically-derived surviving megaspores after meiosis is completed. Following megasporogenesis, extra-numerary cells acquire FM identity but only one of them undergoes gametogenesis to form a female gametophyte containing supernumerary nuclei that do not undergo cellularization or differentiation. The expression of three target genes of miR822a – *At5g02350, At5g02330* and *At2g13900* – is significantly increased in *mir882-1* and *ago9* mutants, and their overexpression in wild-type lines result in defects equivalent to those found in mutant *mir822-1*. These results suggest that the AGO9-interactor miR822a modulates monosporic development in Arabidopsis, thus revealing for the first time the role of a miRNAs in the regulation of megaspore formation in flowering plants.

## RESULTS

### *MIR822* is specifically expressed in developing ovules

Previous immunoprecipitation results have demonstrated that AGO9 can bind to a selected group of miRNAs (Duran-Figueroa and Vielle-Calzada, 2010, Olmedo-Monfil *et al*., 2010). To determine if some of these miRNAs could have an AGO9-dependent function for female gametogenesis, we analyzed the spatial and temporal expression pattern of seven AGO9-interacting miRNAs during flower development. We performed transcriptional fusions between the transcription regulatory region of different *MIR* genes and the *uid*A (GUS) reporter gene. For each regulatory region, we selected at least 500 nt upstream of the first nucleotide at the 5’-end of the predicted miRNA precursor (Zhou *et al*., 2007), and analyzed the GUS expression pattern in at least three independently transformed lines for two consecutive generations. We observed that the promoters from *MIR390* and *MIR161* drove expression of *uidA* in all reproductive organs throughout development (Figure 1); however, while *MIR161pro:GUS* lines did not show expression in the anther, *MIR390pro:GUS* lines showed expression in all regions of male and female reproductive organs. The promoter of *MIR858* only showed expression in the receptacle and sepals (Figure 1), whereas the regulatory regions from *MIR867, MIR403* and *MIR158* drove expression of *uidA* only in the anthers (Figure 1). Whereas *MIR867pro:GUS and MIR403pro:GUS* lines only showed expression during floral stages 10 to 12 and 11-12 respectively, *MIR161pro:GUS, MIR390pro:GUS, MIR858pro:GUS* and *MIR158pro:GUS* lines maintained expression throughout flower development. By contrast, *MIR822pro:GUS* lines showed specific expression in the developing ovules, the expression of the reporter GUS in the ovule initiated at floral stage 11, corresponding to early stages of megasporogenesis, and remained throughout flower development.

Having observed that the expression of GUS in the *MIR822pro:GUS* lines occurred in the ovules of the developing flower (Figure 2A), we next sought to follow the expression of the reporter throughout ovule development. As illustrated in Figure 2, we saw that reporter expression in *MIR822pro:GUS* lines is absent from the MMC at early stages of differentiation (Figure 2B), suggesting that *MIR822* is not transcriptionally active during MMC specification. However, during meiosis, reporter expression starts to be observed in cells of the young inner integuments (Figure 2C) and, at the end of megasporogenesis GUS expression expands to the entire distal pole of the ovule (micropylar region), including in the developing inner and outer integuments (Figure 2D and 2E), suggesting that sporophytic cells actively participate in the transcription of the *MIR822* gene. During female gametogenesis, GUS expression is observed in the micropylar region of the ovule and in the *syncytium* structure, including gametic cells (Figures 2F and 2G). In addition, *MIR822pro:GUS* lines showed reporter expression in trichomes and seedlings, but not in anthers or seeds throughout development. These results indicate that the expression of the *MIR822* initiates at the onset of meiosis during megasporogenesis, and that *MIR822* is active in sporophytic and gametic cells of the ovule.

**Figure 1.**
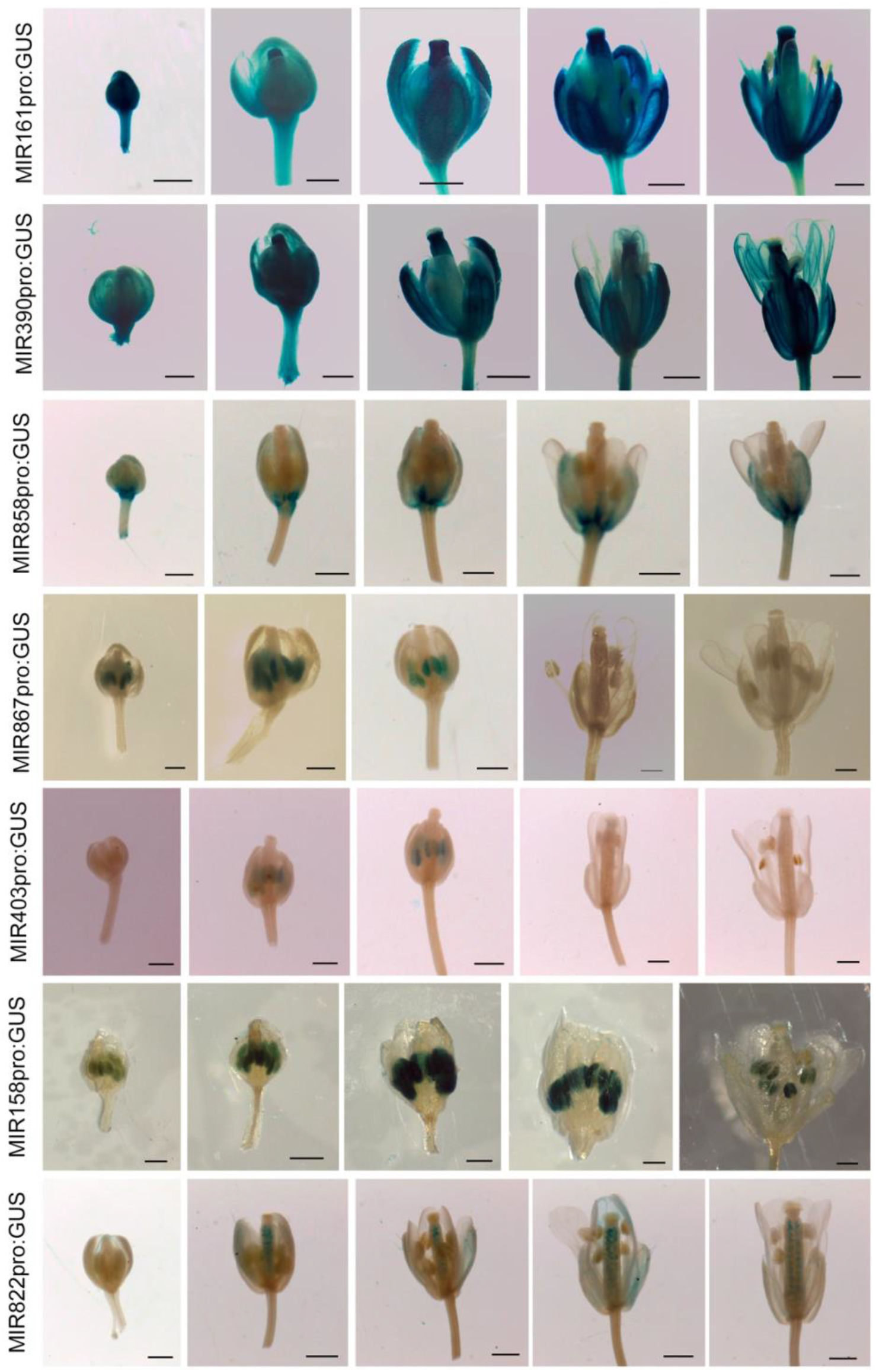
Expression pattern of MIR genes during flower development

**Figure 2.**
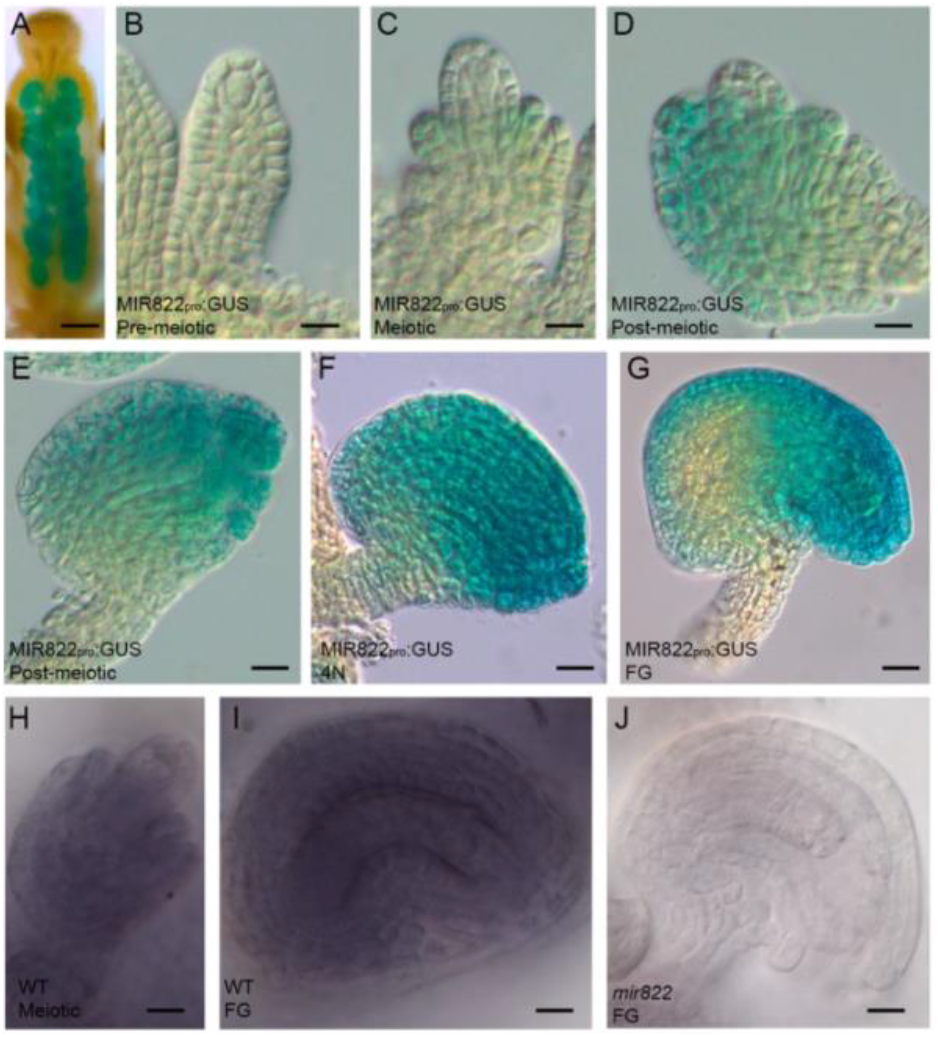
miR822a has a specific expression pattern during ovule development.

To determine the spatial-temporal expression pattern of the mature miR822a, we conducted *in situ* hybridization experiments in wild-type and mutant *mir822* plants. First, we quantitatively assessed the level of expression of miR822a in wild-type plants and in two insertional mutant lines by qRT-PCR. One of the mutant lines (Salk_023928, subsequently named *mir822-1*) has a T-DNA inserted in nucleotide 77 towards the 3’of the *MIR822* gene, and the other one (Sail_99_F11, subsequently named *mir822-2*) has a T-DNA inserted within the predicted *MIR822* promoter, 613 nt upstream of the 5’
send of miR822a (Supplemental Figure 2). Whereas wild-type plants showed up to 4-fold change in the expression of miR822a in ovules, gynoecia, and whole mature flowers, ovules from homozygous *mir822-1* plants showed null expression of the mature miR822a. Conventional RT-PCR using specific primers to detect the *MIR822* precursor, confirmed that the corresponding non-coding RNA is absent in the ovules of *mir822-1*. Then, *in situ* hybridization confirmed that the mature miR822a is localized in the wild-type ovules prior to meiosis and throughout megagametogesis (Figure 2H and 2I) including gametic cells, but absent from mutant *mir822-1* ovules (Figure 2J). Overall, these results confirm the presence of the mature miR822a during megaspore formation and indicate that *miR822-1* is a mutant allele with no expression of miR822a.

### Ovules impaired in *MIR822* function show more than one surviving megaspore

To explore de function of miR822a during female gametogenesis, we investigated if mutant alleles *mir822-1* and *mir822-2* showed any sign of fertility defects. Thus, we performed quantitative morphological analysis of young seeds in open siliques and found that homozygous individuals of both *mir822-1* and *mir822-2* showed a semi-sterile phenotype, with 16.78% and 18.36%, respectively, of unfertilized ovules aborting before seed maturity (Figure 3A-3C). To determine the cellular defect that causes this semi-sterile phenotype, we compared ovule development in wild-type and mutants individuals using whole-mounted cleared specimens under bright field or confocal microscopy, following previously defined megasporogenetic stages on the basis of integument formation (Rodriguez-Leal *et al*., 2015). As expected, in most cases wild type (94.4%; n=180) ovules differentiated a single MMC that then underwent meiosis to give rise to a unique functional megaspore (Figure 3D-3E). By contrast, although most homozygous *mir882-1* and *mir822-2* ovules also differentiated a single pre-meiotic MMC (Figure 3F) (97.5% and 98.07%, respectively; n=160 and n=260), both mutant alleles showed a high frequency of supernumerary derived cells aligning in the micropylar-chalzal axis orientation and resembling non-degenerated meiotic products (Figure 3G to 3J). These supernumerary cells were present from the early stages of integument formation up to stages in which the outer integument is fully developed and surrounding the nucellus. In the case of *mir822-1*, 27.69% of developing ovules showed one additional cell while 7.04% showed two additional cells (n=213). In the case of *mir822-2*, 25.38% showed one additional cell and 6.73% showed two additional cells. Because miR822 is an interactor of AGO9 protein, we made the double mutant *mir822-1 ago9-3* to evaluate the genetic relationship. We observed that in heterozygous plants 33.02% of developing ovules showed one additional cell (Figure 3K). To determine if supernumerary cells are indeed derived from meiosis and acquire a gametic identity, we monitored the expression of *DISRUPTION OF MEIOTIC CONTROL1* (*AtDMC1*) and *CIHUATEOTL* (*CIH*; At4g38150) in ovules of the *mir822-1* mutant. Whereas the *AtDMC1* gene is essential for homologous recombination in meiosis and its expression is located in the MMC (Klimyuk and Jones, 1997, Seeliger *et al*., 2012), *CIH* encodes a pentatricopeptide-repeat protein that is specifically expressed in the FM at the onset of female gametogenesis but, not in the meiotically-derived degenerated megaspores (Sanchez-Leon *et al*., 2012). We independently crossed transgenic lines carrying a transcriptional reporter fusion of the AtDMC1 or pFM2 promoter (pAtDMC1:GUS and pFM2:GUS, respectively) to homozygous *mir822-1* individuals, and histochemically analyzed. Mutant *mir822-1* ovules showed restricted expression of the pAtDMC1:GUS marker in the MMC, confirming a correct onset of meiosis (Figure 3L and 3M). Then, to demonstrate if meiotic division were properly carried out, we monitored the callose deposition with aniline-blue fluorescence (Rodkiewicz, 1970). We confirmed that in ovules of mutant *mir822-1*, callose deposition occurs in a pattern equivalent to wild-type during meiosis I, the callose is deposited in the transverse wall of the newly synthesized cell plate (Figures 3N and 3O). By contrast, following meiosis II, the accumulation of callose in dying megaspores did not occur in *mir822-1* ovules that only showed callose deposition in cell walls transversally oriented on the basis of the micropylar-chalazal axis (Figure 3P and 3Q). The absence of callose deposition in FM adjacent cells is indicative of megaspore survival following meiosis II. Finally, homozygous *mir822-1* ovules showed pFM2 driven GUS expression in two adjacent post-meiotic cells, (Figure 3R and 4S), confirming thus that the abnormally surviving cell acquire the identity of a functional megaspore and are of meiotic origin. Taken together, these results indicate that *MIR822* is necessary for restricting the survival of meiotically derived cells to a single functional megaspore. Also, suggest that *MIR822* with *AGO9* modulate the monosporic female development.

**Figure 3.**
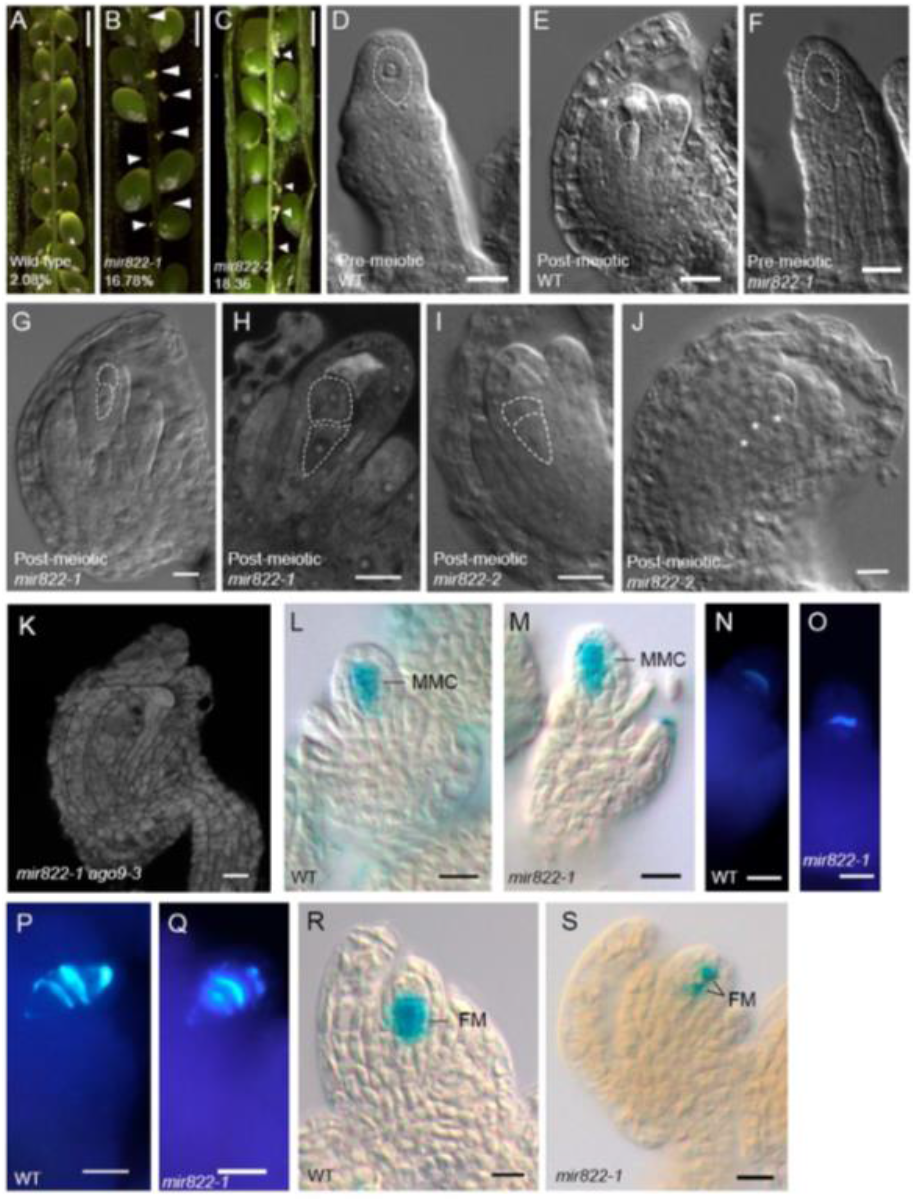
Phenotype of *mir822a* during female gametophyte development

**Figure 4.**
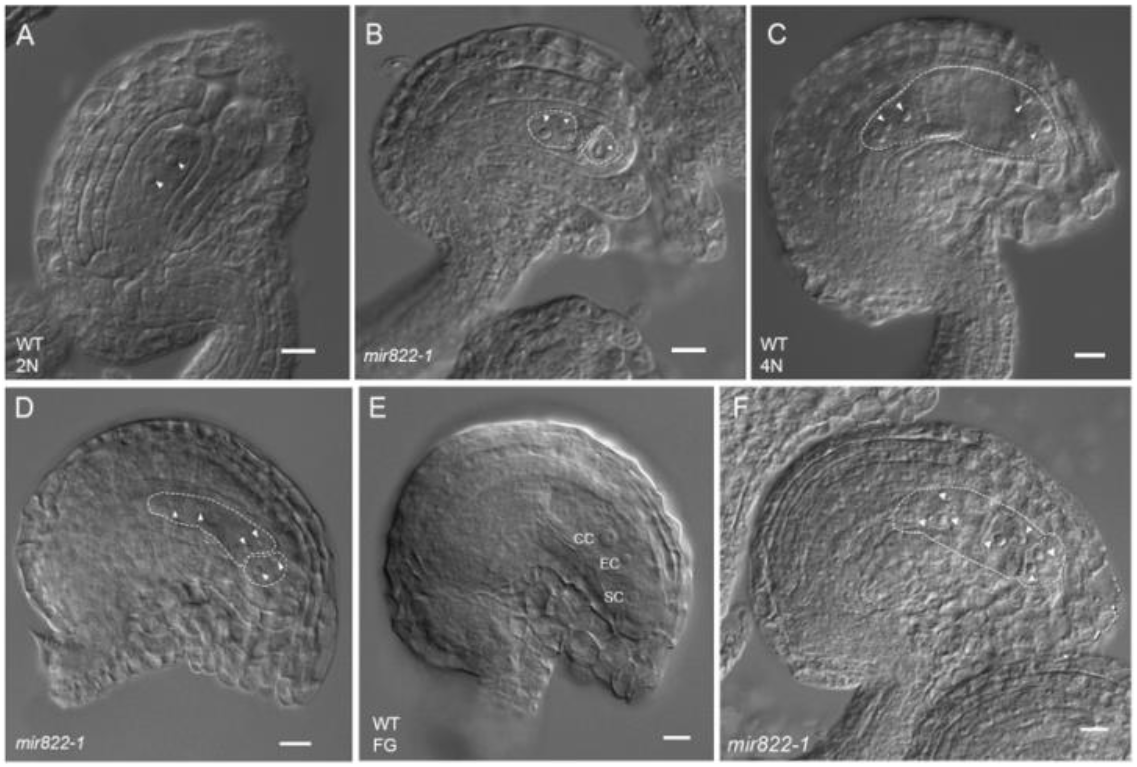
Female megagametogenesis in wild-type and mir822 mutant ovules

To determine if supernumerary FM-like cell is able to subsequently divide and if a female gametophyte is formed, we cytologically compared female gametogenesis in wild-type and *mir822* mutant ovules. As in the case of wild-type, homozygous *mir822-1* and *mir822-2* individuals showed a single FM and subsequently a two nuclear (2N) vacuolated female gametophyte occupying the predicted position, indicating that at this stage of development the abnormally surviving FM-like cell have not initiated female gametogenesis (Figure 4A and 4B). However, once the normally dividing female gametophyte reached the four nuclear stage (4N), 31.62% (n=117) of *mir822-1* and 23.07% (n=117) of *mir822-2* ovules showed an additional 2 nuclear female gametophyte that did not vacuolate or expand (Figure 4C and 4D). Finally, at stages in which a fully differentiated female gametophyte is present in the mature ovule (Figure 4E), *mir822-1* plants had 28.92% (n=242) of ovules showing multinuclear female gametophyte with no apparent cellularization of gametophytic cells (Figure 4F); in the case of *mir822-2*, this frequency was of 26.15% (n=195). These results indicate that in the absence of *MIR822* function, abnormal surviving megaspores are able to mitotically divide but, not to form a cellularized female gametophyte.

### miR822a target genes are regulated by an AGO9-dependent pathway and his overexpression results in gametophytic defects equivalent to those found in *mir822* plants

To confirm that the previously described mutant phenotype is caused by the absence of the miR822a regulatory role during megasporogenesis, we quantitively assessed the expression of three miR822a target genes and determined their function during early ovule development. A combination of high-throughput deep sequencing and RNA ligase-mediated 5′ Rapid Amplification of cDNA ends assays (RLM 5′ RACE), has previously showed that three genes encoding Cysteine/Histidine-Rich C1 domain proteins of unknown function - *At2g13900, At5g02330*, and *At5g02350* – are the targets of miR822a (Addo-Quaye *et al*., 2008). As illustrated in Figure 5A, qRT-PCR assays showed that all three genes are overexpressed in homozygous *mir822-1* ovules as compared to wild-type, confirming their direct regulation by miR822a as previously suggested by (Shao *et al*., 2013). Although it is well known that AGO9 protein preferentially binds 24 nt siRNAs, immunoprecipitation assays had shown that that miR822a is also an AGO9 interactor (Duran-Figueroa and Vielle-Calzada, 2010, Olmedo-Monfil *et al*., 2010). To determine whether AGO9 regulates the expression of these three miR822a target genes, we also assessed their expression by qRT-PCR in ovules of *ago9-3* homozygous individuals. All three showed high levels of expression as compared to wild-type ovules (Figure 5B), reaching up to 1000-fold differences in the case of *At2g13900*. These results suggest that miR822a acts through an AGO9-dependent pathway to negatively regulate Cysteine/Histidine-Rich C1 domain proteins in the ovule.

**Figure 5.**
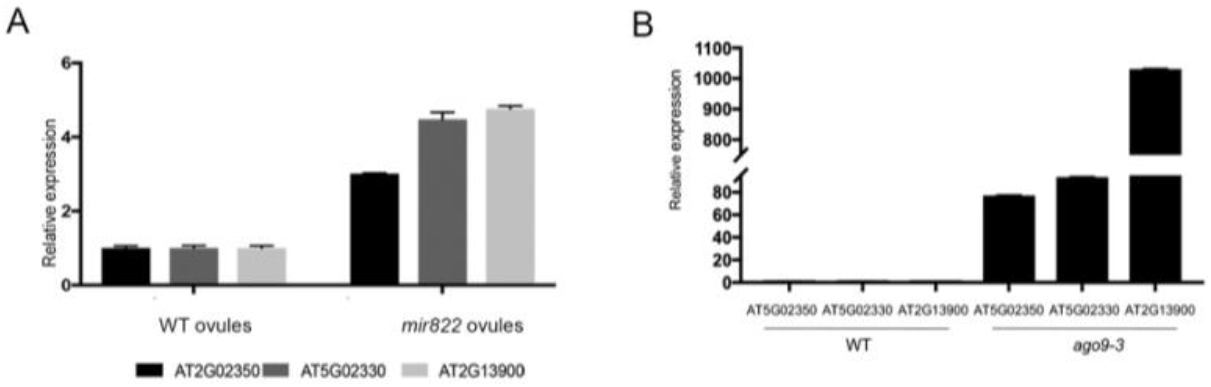
The genes AT5G02350, AT5G02330 and AT2G13900 are downregulated by miR822 and AGO9.

To determine if repression of *At2g13900, At5g02330* and *At5g02350* is necessary to ensure the survival of a single megaspore following meiosis, we generated independent transgenic lines overexpressing each of these three genes under the control of the cauliflower mosaic virus 35S promoter (CaMV35S). Previous results have shown that the CaMV35S promoter is useful to elucidate the function of genes acting during megasporogenesis (Demesa-Arevalo and Vielle-Calzada, 2013). After the generation of three overexpression lines for each target gene, we selected the transgenic lines with the highest level of overexpression to perform a phenotypic analysis (Supplementary Figure 3). Similar to *mir822-1* and *mir822-2*, all overexpressing lines showed a semi-sterile phenotype, with 18.17% (*At5g02350*), 19.33% (*At5g02330*) and 14.6% (*At2g13900*) of the unfertilized ovules aborting before seed formation (Figure 6A). In all cases, developing ovules did not show cytological defects prior to meiosis, suggesting that none of these genes has a role in MMC specification (Figure 6B). By contrast, abnormally surviving megaspores are present when any of these three genes is overexpressed in the Arabidopsis ovule. Whereas the overexpression of *At2g13900* resulted in 36.5% (n=82) of ovules showing an additional surviving megaspore, overexpression of *At5g02330* and *At5g02350* resulted in the same phenotype at frequencies of 31.6% (n=79) and 31.1% (n=72), respectively. These results demonstrate that the repression of any of these three miR822a target genes is necessary to maintain monospory in Arabidopsis, confirming that this AGO9-dependent miRNA regulatory pathway is necessary to inactivate Cysteine/Histidine-Rich C1 domain proteins in the ovule, and restrict the survival of meiotically derived cells to a single megaspore.

**Figure 6.**
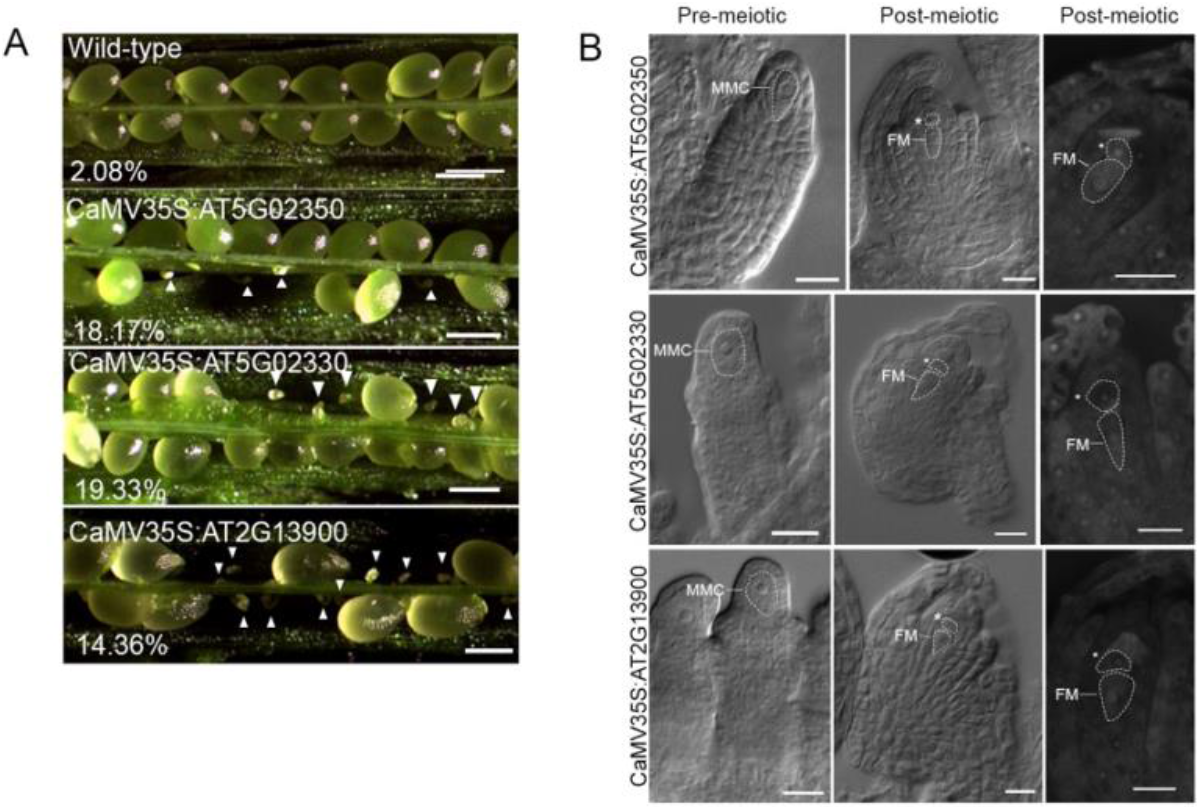
Overexpression of genes *AT5G02350, AT5G02330* and *AT2G13900* phenocopy the *mir822*.

## DISCUSSION

In this paper we have shown that a microRNA modulates monosporic female gametogenesis through an ARGONAUTE9-dependent pathway in *Arabidopsis thaliana*. We have shown that AGO9-miRNAs interactors have a particular temporal and spatial expression pattern during flower development but, evidence suggest that *MIR822* play a key role during early ovule development. We carried out a cellular and molecular characterization of the *loss-of-function* mutant of *MIR822* and showed that extranumerary cells survive towards the end of meiosis instead of one, as it occurs in monosporic development of wild-type plants. By monitoring the AtDMCI::GUS and callose molecular markers in the *mir822-1* mutant background, we demonstrated that surviving-cells are products of meiosis but also interestingly, cells acquire FM identity, which we confirmed with the pFM2 molecular marker. Previously reported degradome and bioinformatic studies, have shown that miR822a has *AT2G13900, AT5G02330* and *AT5G02350* as target genes. We showed that these three genes are overexpressed in the absence of either miR822a or AGO9 protein, indicating that the complex AGO9-miR822a negatively regulates the expression of these three genes during ovule development. Not surprisingly, the overexpression of these three target genes yielded lines that had a phenotype undistinguishable from the *mir822-1* and *mir822-2* alleles during megasporogenesis. Taken together, our data suggest that an AGO9-miR822a pathway modulates the monosporic development by restrict the survival of meiotically derived cells to a single megaspore during megasporogenesis in *Arabidopsis thaliana*.

Although the understanding of the mechanisms controlling megasporogenesis are limited, some new insights have emerged in recent years. Genes controlling early steps of germline fate and meiosis have been identified by several mutant screens (reviewed by (Erbasol Serbes *et al*., 2019)). A substantial amount of data exists that support our current understanding on restriction of MMC fate, and to a certain extent also about the process governing meiotic division of the MMC (Pinto *et al*., 2019). However, much less is known about the mechanisms that ultimately led to the selection of a FM through the degeneration of three and survival of one megaspore. Questions such as *how is monosporic development controlled?* or, *how is megaspore death or survival controlled?* remain largely unanswered. The formation and maturation of the initial cells deriving from meiosis gives rise to the megaspores, depending on the pattern of cell wall formation, megasporogenesis is classified into three types based on the number of nuclei incorporated in the resulting female gametophyte (Rodkiewicz, 1970). The *Polygonum* type of monosporic development prevails in more than 70% of flowering plants – including Arabidopsis – and depends on a single haploid precursor, usually located at the chalazal pole, giving rise to the female gametophyte. According to theoretical hypothesis of genetic conflict during megasporoegensis, the mosoporic type of development in the most stable form of female gametogenesis, ensuring the absence of direct competition among megaspores (Haig, 1990, Bachelier and Friedman, 2011). In some plant families, the surviving megaspore is located on the distal pole (*Oenothera*), which suggests that there are positional signals and therefore, there are a spatial determination of megaspores. The plasticity of developmental patterns observed in angiosperms has allowed to postulate that a failure in cell division and cytokinesis, can lead to the survival of two or four megaspores, which correspond to a bisporic (*Allium*, 8-nucleate) or tretrasporic (*Drusa*, 16-nucelate) development, respectively (Huang and Russell, 1992). In monosporic development of *Arabidopsis thaliana*, the cytokinesis occurs only after completion of meiosis, sometimes giving rise to a multiplanar tetrad of megaspores (Webb and Gunning, 1990, Schneitz *et al*., 1995, Bajon *et al*., 1999). The mechanisms that regulate cytokinesis during female gametogenesis are largely unknown. In other hand, cell-cycle regulators such as genes encoding interactor/inhibitor of cyclin-dependent kinase/Kip-related proteins (*ICK/KRPs*) have been associated to the degeneration of megaspores (Cao *et al*., 2018). Other kind of genes located in plasma membrane, that include *ARABINOGALACTAN PROTEIN18* and *ARABIDOPSIS HISTIDINE KINASE* have been associated to FM selection (Cheng *et al*., 2013, Demesa-Arevalo and Vielle-Calzada, 2013); but their role in degradation or survival is nor clearly understood.

Here we show that a miRNA is involved in FM selection at the end of megasporogenesis. Our results shown that in the *mir822* mutant and the overexpressing lines of the corresponding target genes, have a bisporic-like development at high frequency. Interestingly, the presence of extra-numerary megaspores in mutant and overexpresses lines did not give rise to an additional, differentiated and functional embryonic sac, indicating, perhaps not surprisingly, the existence of other regulatory elements controlling the stages of female gametophyte development. Remarkably, the surviving extra FM-like cell is derived from meiosis II and invariably is the cell adjacent to proximal position, therefore suggesting that elimination of cells is being modulated by a positional signal. Together with programmed cell death (PCD), positional signals are a developmental strategy in seed formation to control cell number and identity (Ingram, 2017). Nevertheless, genes associated with this cellular process and with implications in megasporogenesis have not been identified to date. We postulate that, miR822a mediates a pathway involved in the cell-elimination of haploid products after meiosis during megasporogenesis. It is possible then, that miR822a works like an intercellular signal for triggering PCD during megasporogenesis. Already some decades ago, it was documented that one hallmarks of megaspores degeneration is the deposition of callose since this accumulates positionally in those cells destined to die (Rodkiewicz, 1970, Kapil and Bhatnagar, 1981). In the mutant background of *mir822*, callose signal is clearly observed after cell division in meiosis I and meiosis II, just as in the wild-type case, but the typical callose deposition towards the distal pole is not observed in the mutant. Lack of callose has been suggested with the free flow of small RNAs in the plasmodesmata of the haploid megaspores (Tucker and Koltunow, 2014). Our evidence suggests that the function of miR822a is to induce cell elimination of female haploid products and thus, allows only one cell to survive to develop the mature and functional embryo sac in a monosporic type development. Interestingly, the role of miR822a in monosporic development might be restricted to the Arabidopsis genera, because *MIR822* gene is a non-conserved MIR gene (Chavez Montes *et al*., 2014).

Although more detailed experiments are necessary, the combination of transcriptional fusions experiments and *in situ* assay indicates that mature miR822a is located in both sporophytic and gametophytic cells but, its synthesis apparently is restricted to sporophytic cells, resembling a non-cell autonomous mechanism (Nonomura, 2018). It has been widely demonstrated that miRNAs and siRNAs can move over long distances as a mechanism of cellular communication, move also between cells and act as positional signals (Benkovics and Timmermans, 2014, Liu and Chen, 2018). Our previous studies it has been suggested that TE-derived siRNAs that are loaded by AGO9 restrict the cellular identity of MMC by a non-cell autonomous pathway (Duran-Figueroa and Vielle-Calzada, 2010, Olmedo-Monfil *et al*., 2010). Furthermore, it has been shown that a pathway mediated by a complex siRNA/AGO can promote meiotic division of megaspores (Tucker *et al*., 2012) and, also has been suggested that a sporophytic source of cytokinin is required for correct FM specification (Cheng *et al*., 2013). Therefore, it remains to be clarified if the miR822 moves, intercellularly from young integuments to gamete precursor cells during megasporogenesis and if works as positional signal for modulating the monosporic developmental pattern in Arabidopsis.

As we have also shown in this paper, expression of, miR403, miR867 or miR57, which are all AGO9 interactors, is restricted to pollen, opening up a great opportunity to study the role of the AGO9-miRNA complexes in plant male organ development. Evidence that AGOs works in nucleus and cytoplasm has been documented (Bologna *et al*., 2018). Previously published results by our group have shown that AGO9 is located in both the nucleus and the cytoplasm, particularly in pollen it is localized in cytoplasmic foci of the vegetative cell (Olmedo-Monfil *et al*., 2010, Rodriguez-Leal *et al*., 2015). It is important to highlight that the miRNAs evaluated in this work are also interactors of AGO1 (Havecker *et al*., 2010); however, neither the AGO1 nor the genes associated with miRNA biogenesis have been associated with megasporogenesis (Hernandez-Lagana *et al*., 2016).

Finally, an open question to investigate in the future is: *which is the actual function of target genes regulated by miR822a?* The three target genes *AT5G02350, AT5G02330 and AT2G13900* belong to a huge family of approximately 140 genes in Arabidopsis. These genes contain cysteine-and histidine-rich zinc finger domain called Divergent C1 (DC1) and, are found exclusively in plants. The DC1 domain is still poorly understood, few reports have approached them to characterize it and study its role. The DC1-domain gene called *VACUOLELESS GAMETOPHYTES* (*VLG*) has been characterized in Arabidopsis and shown to be necessary for vacuole formation at early development in both male and female gametophytes (D’Ippolito *et al*., 2017). It is very interesting that in the *mir822* mutant, the classical vacuole that forms at the 2N stage of development is absent. In *Capsicum annuum*, a gene coding for a protein with DC1 domains has been shown to be involved in salicylic acid dependent plant defense response and, has also been associated with plant cell death; additional experiments showed that this protein is capable of binding both DNA and RNA *in vitro* (Hwang *et al*., 2014). In cotton, a DC1 domain-containing transcription factor has been identified as the target of miRNVL5 and the latter as regulator of the plant response to salt stress (Gao *et al*., 2016). It would be very interesting to determine whether the genes *AT5G02350, AT5G02330* and *AT2G13900* are transcription factors involved in vacuole formation with consequences on PCD associated to female gametogenesis in *Arabidopsis thaliana*.

## MATERIALS AND METHODS

### Plant Material and culture conditions

Wild-type *Arabidopsis thaliana* Columbia ecotype (Col-0) was used for all experiment involving transformation. Seeds were surface sterilized in a solution of 60% (v/v) commercial chlorine and 0.005% Tween 20 (Sigma, USA). Seeds were placed in Murashige and Skoog (MS) (Murashige and Skoog, 1962) medium and incubated for germination in a chamber with long-day (16 h light/8 h dark) conditions at 25ºC. Selection of transformed lines was carried out in MS medium supplemented with kanamycin (50 mg/mL) and using the same environmental conditions already mentioned. For seed collection, plants were grown under greenhouse conditions at 25ºC. All transgenics plants selected were in the F3 generation. The *mir822-1* (*Salk_023928)* and the mir822-2 (*Sail_99_F11*) lines were genotyped following the Salk Institute Genomic Analysis Laboratory instructions, and absence of the pre-miR822 was confirmed by end-point RT-PCR. The *mir822*x*pAtDMC1-GUS* and *mir822*x*pFM2-GUS* were obtained crossing the homozygous Salk_023928 line with each marker line. All primers used for genotyping are listed in Supplementary Table 1.

### Vectors Construction and transformants generation

The core of the Promoter Region (PR) from MIR genes was selected considering genomic characteristic according to Zhou and collaborators (Zhou *et al*., 2007). Each PR, was amplified by PCR from wild-type genomic DNA (Primers used are listed in Supplementary Table 1). Amplified PRs had the following lengths: for MIR822 (AT5G03552), 884 bp; for MIR390 (AT2G38325) 985 bp; for MIR858 (AT1G71002) 999 bp; for MIR157 (AT1G66783) 1008 bp; for MIR867 (AT4G21362) 1012 bp; for MIR161 (AT1G48267) 999 bp; and for MIR403 (AT2G47275) 1000 bp. The PRs were independently inserted into pBlueScriptII KS (-) and then transferred using *Hind*III and *Xba*I into binary vector pBI101.3 to generate GUS (*uidA*) fusions. Target genes for miR822, AT5G02330, AT5G02350 and AT2G13900 genes were amplified by PCR from wild-type genomic DNA and cloned into donor vector pENTR/TOPO (Invitrogen). The coding sequences were then transferred to destination vector pGWB2 vector by LR recombination (Invitrogen) to obtain the transformation vectors carrying the expression cassettes CaMV35S:AT5G02330, CaMV35S:AT5G02350 and CaMV35S:AT2G13900. *Arabidopsis thaliana* Col-0 was transformed by the floral dip method (Zhang *et al*., 2006) using *Agrobacterium tumefaciens* strain C58C1. Twenty transformed plants from each line were analyzed by PCR. All cloning and analysis primers used are listed in Supplementary Table 1.

### Expression analysis by qRT-PCR

Total RNA was extracted using ZR Plant RNA MiniPrep (ZymoResearch) kit. For miRNAs, cDNA synthesis was performed with 100 ng of total RNA using miScript Plant RT kit (Qiagen); qRT-PCR assays were done with SYBR Green PCR kit (Qiagen) following the manufacturer’s instructions. LNA probes used for qRT-PCR to detect mature miRNAs were: miScript Primer Assay At_miR822_5p_1 (5’UGCGGGAAGCAUUUGCACAUG) and as a control we used miScript Primer Assay At_mir167_5p_1 (5’UGAAGCUGCCAGCAUGAUCUA). For the target genes AT5G02350, AT5G02330 and AT2G13900 cDNA was synthesized using M-MuLV Reverse Transcriptase (NEB) with 100 ng of total RNA at 42ºC for 60 min, and 65ºC for 20 min for inactivation. The thermal profile of qRT-PCR assays consisted of 95ºC for 10 min, 40 cycles of 95ºC for 15 s, 60ºC for 30 s and 72ºC for 30 s. Each quantitative PCR (qRT-PCR) reaction was performed in a final volume of 10 µL, 5 µL of Maxima SYBR Green/ROX qPCR Master Mix (Thermo Scientific), 0.5 µL of each primer, forward and reverse (10 µM), 6 µL of RNase-free water and 1 µL of cDNA. For ovules we used 50 ng of total RNA for qPCR of target genes. Amplification of *UBIQUITIN* gene was used as control. All reactions were carried out in Eco. Illumina 1010180, and data analyzed using the EcoStudy Illumina software. To determine the relative expression of each gene in different tissues we used the 2^-ΔΔCq^ method. Each qRT-PCR reaction had three biological replicates for all tissues analyzed. Primers used for qRT-PCR assays are listed in Supplementary Table 1.

### Whole-mount *in situ* Hybridization

Whole mount hybridization was based on (Garcia-Aguilar *et al*., 2005) with minor modifications. Inflorescences were fixed in 4% fresh paraformaldehyde in tubes of 1.5 mL and put into desiccator with a vacuum pump for 5 min. Tubes were then placed in continuous agitation for 3 h at room temperature. After fixation, the inflorescences were washed three times with 1X PBS. Inflorescences were rinsed in 50% ethanol, 25% ethanol, and diethyl pyrocarbonate (DEPC)-treated water for 5 min each. After removing all excess water, inflorescences were dissected with a hypodermic syringe and embedded in 15% acrylamide:bysacrilamide (29:1) over polysine slides (Thermo Scientific). Slides were treated with 150 μL of 0.2 M HCl for 20 min at 25°C and subsequently washed three times: in DEPC-treated water, 1X PBS and DEPC-treated water. Samples were digested in 1 µg/mL proteinase K for 30 min at 37°C in a humid environment. Digestion was stopped by washing serially in 1X PBS and 2 mg/mL of glycine for 2 min, samples were subsequently post-fixed in 4% paraformaldehyde for 20 min at 25°C in a humid environment. Slides were transferred into hybridization solution (6X standard saline citrate [SSC], 3% SDS, 50% formamide, and 0.1 mg/mL tRNA) at 55°C for 90 min in agitation. After pre-hybridization LNA probes were added into the hybridization solution and samples were maintained in a humid environment at 55°C for 16 h. After hybridization, slides were washed twice in 0.2X SSC–0.1% SDS at 55°C for 10 min, followed by a wash in 2X SSC for 2 min with gentle agitation. Slides were digested with RNase A (10 µg/mL) in 2X SSC for 30 min at 37°C humid environment. Digestion was stopped by washing with 2X SSC for 2 min at 25°C. Final washes were conducted in 0.2X SSC–0.1% SDS at 55°C for 10 min in agitation, followed by a wash in 2X SSC for 2 min and a wash in 1X TBS (100 mM Tris-HCl [pH 7.5], 150 mM NaCl) for 2 min, both at 25°C with gentle agitation. Slides were incubated in 0.5% blotting-grade blocker (BioRad) for 2 h in TBS and subsequently rinsed in TBS for 2 min at 25°C with gentle agitation. For immunological detection, each slide without coverslip was incubated with 150 µL of anti-digoxigenin–conjugated antibody (Abcam) diluted 1:1000 in 1% bovine serum albumin (BSA) in TBS for 2h at 4°C in a humid environment, followed by 4 washes for 10 min each in 0.5% BSA and 0.1% Triton X-100 with gentle agitation. The slides were subsequently washed in buffer C with levamisole (100 mM Tris [pH 9.5], 50 mM MgCl2, 100 mM NaCl, 0.1% Tween 20, 1 mM levamisole) and incubated for 12-17 h in 0.34 mg/mL nitrotetrazolium blue chloride (Sigma Aldrich) in buffer C with levamisole in a humid environment, each slide was covered with a coverslip. Slides were observed in Normaski optics using a Leica microscope DM5000B.

### Cytological analysis

For DIC microscopy, inflorescences were rinsed and fixed in FAA solution (10% formaldehyde, 5% acetic acid, and 50% ethanol) for 24 h at 4ºC. After fixation, samples were dehydrated in 70% ethanol at room temperature. Gynoecia from different stages of ovule development were dissected with hypodermic needles (BD Plastipack) and observed with a Stereomicroscope Leica EZ4 HD; then, ovules were cleared in Herr’s solution (phenol:chloral hydrate: 85% lactic acid:xylene:clove oil in a 1:1:1:0.5:1 proportion), and observed in Normaski optics using a Leica microscope DM5000B. For MIR822 promoter expression analysis, fresh tissues were incubated in GUS staining solution (10 mM EDTA, 0.1% Triton X-100, 5 mM potassium ferrocyanide, 5 mM potassium ferricyanide, and 1 mg/mL of X-Gluc in 50 mM of phosphate buffer pH 7.4) for 24 h at 37ºC. After incubation, ovules were mounted in a solution of 20% glycerol and 20% of lactic acid. Confocal imaging was performed using Propidium Iodide (PI), inflorescences were fixed in FAA solution and dehydrated in 70% ethanol. Carpels were dissected to obtain ovules and were stained in 1 mg/mL of PI for 6 h at room temperature, ovules were mounted and observed using the Laser Scanning Confocal Microscope (LSCM) Leica TCS SP8.

## ACKNOWLEDGMENTS

Research in the laboratory of NVDF was supported by grant from Consejo Nacional de Ciencia y Tecnologia (CONACyT-184245). Grants 20182227 and 20196284 to NVDF from Instituto Politécnico Nacional through Secretaría de Investigación y Posgrado (SIP-IPN) financed part of this research. We thank Dr. Maria del Carmen Oliver Salvador for all her enthusiasm, motivation and discussion during the project. We also would like to thank all undergraduate students who participated in cloning and growing plants. ATA was financially supported by CONACyT and IPN scholarships during her PhD studies.

